# Gut Microbiome Wellness Index 2 for Enhanced Health Status Prediction from Gut Microbiome Taxonomic Profiles

**DOI:** 10.1101/2023.09.30.560294

**Authors:** Daniel Chang, Vinod K. Gupta, Benjamin Hur, Sergio Cobo-López, Kevin Y. Cunningham, Nam Soo Han, Insuk Lee, Vanessa L. Kronzer, Levi M. Teigen, Lioudmila V. Karnatovskaia, Erin E. Longbrake, John M. Davis, Heidi Nelson, Jaeyun Sung

## Abstract

Recent advancements in human gut microbiome research have revealed its crucial role in shaping innovative predictive healthcare applications. We introduce Gut Microbiome Wellness Index 2 (GMWI2), an advanced iteration of our original GMWI prototype, designed as a robust, disease-agnostic health status indicator based on gut microbiome taxonomic profiles. Our analysis involved pooling existing 8069 stool shotgun metagenome data across a global demographic landscape to effectively capture biological signals linking gut taxonomies to health. GMWI2 achieves a cross-validation balanced accuracy of 80% in distinguishing healthy (no disease) from non-healthy (diseased) individuals and surpasses 90% accuracy for samples with higher confidence (i.e., outside the “reject option”). The enhanced classification accuracy of GMWI2 outperforms both the original GMWI model and traditional species-level α-diversity indices, suggesting a more reliable tool for differentiating between healthy and non-healthy phenotypes using gut microbiome data. Furthermore, by reevaluating and reinterpreting previously published data, GMWI2 provides fresh insights into the established understanding of how diet, antibiotic exposure, and fecal microbiota transplantation influence gut health. Looking ahead, GMWI2 represents a timely pivotal tool for evaluating health based on an individual’s unique gut microbial composition, paving the way for the early screening of adverse gut health shifts. GMWI2 is offered as an open-source command-line tool, ensuring it is both accessible to and adaptable for researchers interested in the translational applications of human gut microbiome science.

## Introduction

Recent landmark studies have unveiled profound links between the gut microbiome and a variety of complex chronic diseases^1–9^. Despite these advances, a central challenge remains: how can we harness these unique microbial signatures to develop robust, gut microbiome-centric quantitative assays for non-invasive tracking of human health? This pivotal question represents an essential frontier in utilizing gut microbiota as precise indicators of health and wellness.

The potential of the gut microbiome as a marker for deciphering complex, chronic diseases has captivated the scientific community—in response, we recently developed the Gut Microbiome Wellness Index (GMWI) [previously called the Gut Microbiome Health Index (GMHI)]^10^. GMWI is a first-of-its-kind stool metagenome-based indicator for assessing health by determining the likelihood of an individual harboring a clinically diagnosed disease solely from their gut microbiome state, irrespective of the specific disease type^10,11^. This disease-agnostic index was derived from a comprehensive analysis of a pooled dataset comprising 4347 stool shotgun metagenomes from 34 independent studies. The GMWI is a logarithmic ratio of the collective abundances—a term encompassing species-level relative abundances and multiple α-diversity metrics—of health- and disease-associated gut microbial species. Evaluating on the pooled dataset, GMWI exhibited a balanced accuracy (i.e., average of the proportions of healthy and non-healthy samples that were correctly classified) of 69.7% in predicting the presence of clinically diagnosed disease. Specifically, the correct classification rates for healthy (disease-free) individuals and those with non-healthy (disease-harboring) conditions were 75.6% and 63.8%, respectively. Moreover, the GMWI achieved a balanced accuracy of 73.7% in the validation cohort of 679 stool metagenomes, with the correct classification rates for the healthy and non-healthy subsets being 77.1% (91 out of 118) and 70.2% (394 out of 561), respectively. Since its original publication in 2020, GMWI has been utilized in studies investigating the impact of environmental^12^ and genetic/socioeconomic^13^ aspects on the human gut microbiome, and helped identify a ‘Longevous Gut Microbiota Signature’ species set by Xu *et al*.^13^

Despite the promise of our GMWI prototype, there are inherent limitations that impede its general applicability. Firstly, GMWI correctly classifies healthy stool metagenomes at a higher success rate than non-healthy ones. This bias in GMWI’s predictive performance against non-healthy individuals may be attributed to the prevalence-based strategy used to identify health-associated and disease-associated species, which was a fundamental component of our GMWI model. As the non-healthy group encompasses patients with different diseases, this group is inherently heterogeneous; so a prevalence-based strategy may fail to identify subtle taxonomic signatures that are only represented in subsets of non-healthy populations (e.g., cohorts of a particular disease). It is essential to refine this approach to enhance the predictive accuracy of GMWI in non-healthy individuals. Secondly, our existing model assigns equal weight to each species, not considering potential variances in the importance of individual species. To enhance the classification accuracy and general applicability of GMWI, it is necessary to develop a refined weighting system that accounts for varying strengths of association and class distributions. Additionally, incorporating data from all taxonomic ranks allows microbial signatures across various phylogenetic depths to be captured, and may be critical for optimally predicting host phenotype from microbial communities^14^. In this study, we present GMWI2, an enhanced version of the original GMWI that addresses the above limitations, and shows significant advances in classification accuracy in predicting both healthy and non-healthy phenotypes.

## Results

### Pooled analysis of stool metagenomes across health and disease phenotypes

As in our previous work, we define “healthy” subjects as those without reported diseases or abnormal body weight conditions (i.e., classified as underweight, overweight, or obese based on reported BMI), while “non-healthy” are those with a clinical diagnosis of any disease agnostic to a specific phenotype. We conducted a pooled analysis of existing 8069 stool shotgun metagenomes (5547 from healthy individuals and 2522 from non-healthy individuals) sourced from 54 independently published studies (**Fig. 1a**, **Table 1**, and **Supplementary Data 1**). These pooled metagenomes are from individuals with 12 different health and disease phenotypes (**Fig. 1a**; healthy, ankylosing spondylitis, atherosclerotic cardiovascular disease, colorectal cancer, Crohn’s disease, Graves’ disease, liver cirrhosis, multiple sclerosis, nonalcoholic fatty liver disease (or also known as metabolic dysfunction-associated steatotic liver disease [MASLD]), rheumatoid arthritis, type 2 diabetes, and ulcerative colitis) from diverse geographies, ethnicities/races, cultures, and balanced sex representation (**Fig. 1b**). (Our study and sample selection criteria can be found in “Methods”. Additionally, we provide all subjects’ phenotype, age, sex, BMI, and geography [as provided in their respective original study] in **Supplementary Data 2**.) This substantial increase in sample size, nearly doubling the number of metagenomes included in our original study, is one notable improvement in GMWI2. By increasing the diversity and size of the dataset, we can promote broader representation in microbiome research, and enhance the generalizability and applicability of the model to a wider population. Additionally, GMWI2 uses MetaPhlAn3^15^ instead of MetaPhlAn2^16^ for taxonomic profiling, leveraging an extensively expanded marker database for a more comprehensive and accurate characterization of microbial taxa (“Methods”).

**Figure 1.**
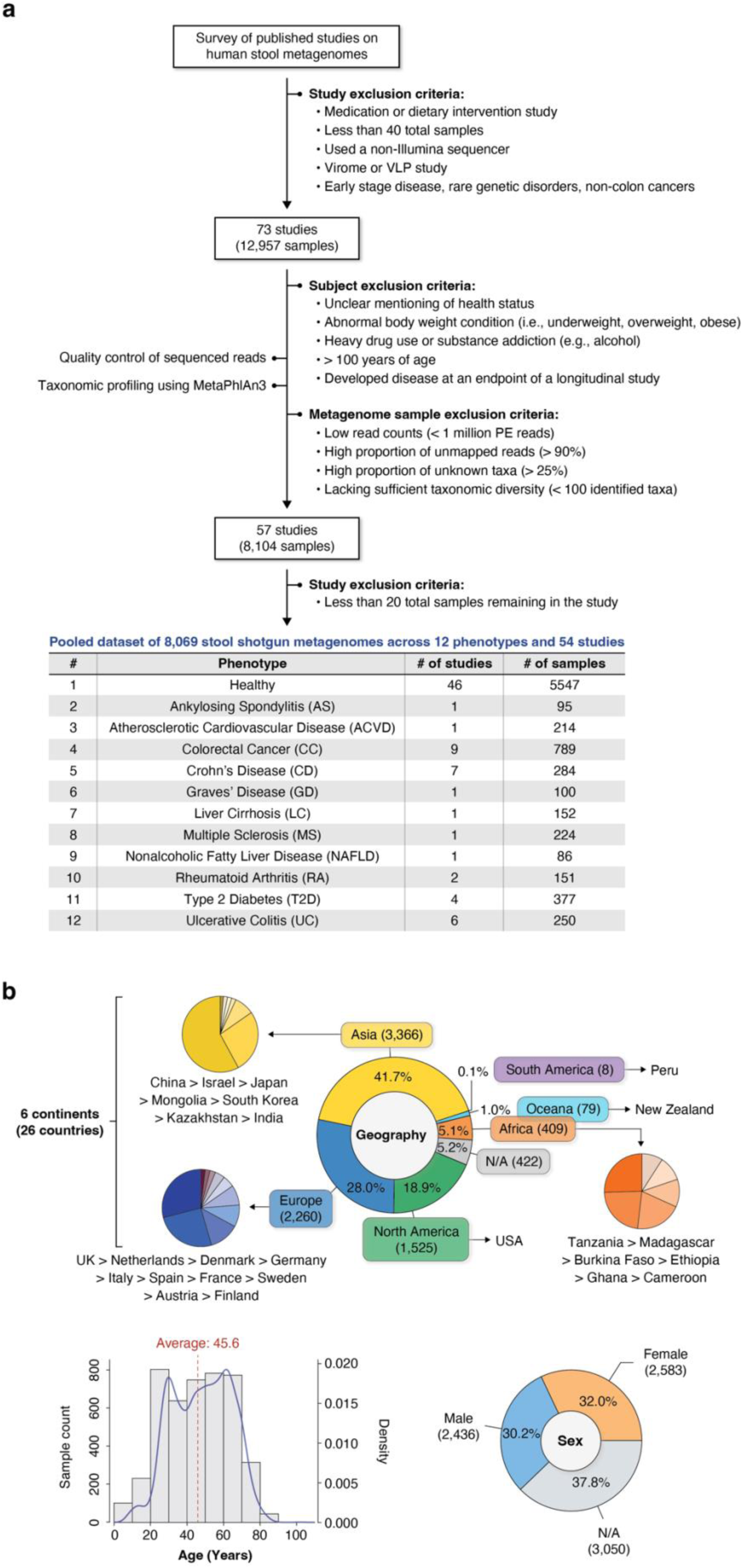
Pooled analysis of stool metagenomes across multiple health and disease conditions from a diverse global representation. **(a)** A survey was conducted in PubMed and Google Scholar to search for published studies with publicly available human stool shotgun metagenome (gut microbiome) samples from healthy (disease-free) and non-healthy (diseased) individuals. The initial collection of stool metagenomes consisted of 12,957 samples from 73 independent studies. All samples (.fastq files) were downloaded and reprocessed uniformly using identical bioinformatics methods. After quality control of sequenced reads, taxonomic profiling was performed using MetaPhlAn3. Studies and metagenome samples were removed based on several exclusion criteria. Finally, a total of 8,069 samples (5,547 and 2,522 metagenomes from healthy and non-healthy individuals, respectively) from 54 studies ranging across healthy and 11 non-healthy phenotypes were assembled into a pooled metagenome dataset for downstream analyses. (**b)** Demographics of the pooled dataset of 8069 human stool metagenomes from 54 published studies. Subject demographics, as reported in the original studies, include age (*n* = 4442), sex (*n* = 5019), and country of origin (*n* = 7647).

**Table 1.** Human stool shotgun metagenome datasets utilized in this study.

| Author<br>(Last name) | Publication<br>year | Total<br>from<br>study (*n*) | Healthy (*n*) | Non-healthy (*n*) | Disease (*n*)<sup>a</sup> | Sequencing platform | Geography (Country) |
| --- | --- | --- | --- | --- | --- | --- | --- |
| Ananthakrishnan | 2017 | 64 | 0 | 64 | CD (24), UC<br>(40) | Illumina NextSeq 500 | United States |
| Ang | 2021 | 22 | 22 | 0 | – | Illumina NovaSeq 6000 | United States |
| Asnicar | 2021 | 568 | 568 | 0 | – | Illumina NovaSeq 6000 | United Kingdom/United States |
| Backhed | 2015 | 100 | 100 | 0 | – | Illumina HiSeq 2000 | Denmark |
| Costea | 2017 | 169 | 169 | 0 | – | Illumina HiSeq 2000 | Germany/Kazakhstan |
| D’Souza | 2021 | 128 | 128 | 0 | – | Illumina NextSeq 500 | Netherlands |
| Davies | 2020 | 44 | 0 | 44 | T2D (44) | Illumina HiSeq 2000 | New Zealand |
| De Filippis | 2019 | 99 | 99 | 0 | – | Illumina HiSeq 1500/Illumina<br>NextSeq 500 | Italy |
| Dhakan | 2019 | 47 | 47 | 0 | – | Illumina NextSeq 500 | India |
| Feng | 2015 | 46 | 0 | 46 | CRC (46) | Illumina HiSeq 2000 | Austria |
| Franzosa | 2018 | 213 | 56 | 157 | CD (84), UC<br>(73) | Illumina HiSeq 2000 | Netherlands/United States |
| Gu | 2017 | 94 | 0 | 94 | T2D (94) | Illumina HiSeq 2500 | China |
| Gupta | 2020 | 49 | 0 | 49 | RA (49) | Illumina HiSeq 4000 | United States |
| He | 2017 | 86 | 40 | 46 | CD (46) | Illumina HiSeq 2000 | China |
| Huttenhower;<br>Lloyd-Price<sup>b</sup> | 2012; 2017 | 507 | 507 | 0 | – | Illumina HiSeq 2000/Illumina<br>Genome Analyzer II | United States |
| Jacobson | 2021 | 82 | 82 | 0 | – | Illumina NovaSeq 6000 | Burkina Faso |
| Jie | 2017 | 322 | 108 | 214 | ACVD (214) | Illumina HiSeq 2000 | China |
| Karlsson | 2013 | 53 | 0 | 53 | T2D (53) | Illumina HiSeq 2000 | Sweden |
| Kim | 2021 | 61 | 61 | 0 | – | Illumina HiSeq 4000 | South Korea |
| Le Chatelier | 2013 | 88 | 88 | 0 | – | Illumina HiSeq 2000/Illumina<br>Genome Analyzer II/Illumina<br>Genome Analyzer IIx | Denmark |
| Liu | 2016 | 110 | 110 | 0 | – | Illumina HiSeq 4000 | China/Mongolia |
| Lloyd-Price | 2019 | 86 | 25 | 61 | CD (39), UC<br>(22) | Illumina HiSeq 2000 | United States |
| Lokmer | 2019 | 37 | 37 | 0 | – | Illumina HiSeq 2000 | Cameroon |
| Loomba | 2017 | 86 | 0 | 86 | NAFLD (86) | Illumina HiSeq 2500 | United States |
| Mehta | 2018 | 301 | 301 | 0 | – | Illumina HiSeq 2000 | United States |
| Nielsen | 2014 | 159 | 82 | 77 | CD (12), UC<br>(65) | Illumina HiSeq 2000/Illumina<br>Genome Analyzer II/Illumina<br>Genome Analyzer IIx | Denmark/Spain |
| Obregon-Tito | 2015 | 20 | 20 | 0 | – | Illumina HiSeq 2500 | Peru/United States |
| Pasolli | 2019 | 142 | 142 | 0 | – | Illumina HiSeq 2500 | Ethiopia/Madagascar |
| Qi | 2019 | 43 | 43 | 0 | – | Illumina HiSeq 2500 | China |
| Qin | 2012 | 369 | 183 | 186 | T2D (186) | Illumina Genome Analyzer II | China |
| Qin | 2014 | 287 | 135 | 152 | LC (152) | Illumina HiSeq 2000 | China |
| Rettedal | 2021 | 35 | 35 | 0 | – | Illumina HiSeq 2500 | New Zealand |
| Roager | 2019 | 50 | 50 | 0 | – | Illumina HiSeq 2000 | Denmark |
| Schirmer | 2016 | 385 | 385 | 0 | – | Illumina HiSeq 2000 | Netherlands |
| Schirmer | 2018 | 83 | 18 | 65 | CD (39), UC<br>(26) | Illumina HiSeq 2000 | United States |
| Smits | 2017 | 38 | 38 | 0 | – | Illumina HiSeq 4000 | Tanzania |
| Sun | 2021 | 42 | 42 | 0 | – | Illumina HiSeq 4000 | United States |
| Tett | 2019 | 110 | 110 | 0 | – | Illumina HiSeq 2000/Illumina<br>HiSeq 2500 | Tanzania/Ghana |
| Thomas | 2019 | 160 | 61 | 99 | CRC (99) | Illumina HiSeq 2500 | Italy/Japan |
| Ventura | 2019 | 48 | 24 | 24 | MS (24) | Illumina HiSeq 4000 | United States |
| Vogtmann | 2016 | 81 | 30 | 51 | CRC (51) | Illumina HiSeq 2000 | United States |
| Wen | 2017 | 200 | 105 | 95 | AS (95) | Illumina HiSeq 2000 | China |
| Weng | 2019 | 79 | 15 | 64 | CD (40), UC<br>(24) | Illumina HiSeq X Ten | China |
| Wirbel | 2019 | 55 | 33 | 22 | CRC (22) | Illumina HiSeq 4000 | Germany |
| Xie | 2016 | 130 | 130 | 0 | – | Illumina HiSeq 2000 | United Kingdom |
| Yachida | 2019 | 217 | 0 | 217 | CRC (217) | Illumina HiSeq 2500 | Japan |
| Yang | 2020 | 180 | 88 | 92 | CRC (92) | Illumina HiSeq X Ten | China |
| Yang | 2021 | 194 | 97 | 97 | CRC (97) | Illumina NovaSeq 6000 | China |
| Yassour | 2018 | 42 | 42 | 0 | – | Illumina HiSeq 2500 | Finland |
| Yu | 2015 | 128 | 53 | 75 | CRC (75) | Illumina HiSeq 2000 | China |
| Zeevi | 2015 | 900 | 900 | 0 | – | Illumina HiSeq 2500/Illumina<br>HiSeq 2500/Illumina MiSeq | Israel |
| Zeller | 2014 | 135 | 45 | 90 | CRC (90) | Illumina HiSeq 2000 | France/Germany |
| Zhang | 2015 | 163 | 61 | 102 | RA (102) | Illumina HiSeq 2000 | China |
| Zhu | 2021 | 132 | 32 | 100 | GD (100) | Illumina HiSeq 4000 | China |

<sup>a</sup>ACVD, atherosclerotic cardiovascular disease; AS, ankylosing spondylitis; CRC, colorectal cancer; CD, Crohn's disease; GD, Graves' disease; LC, liver cirrhosis; MS, multiple sclerosis; NAFLD, nonalcoholic fatty liver disease; RA, rheumatoid arthritis; T2D, type 2 diabetes; UC, ulcerative colitis. <sup>b</sup>Samples combined from both phases of the Human Microbiome Project (HMP1 and HMP1-II).

All metagenomes underwent uniform reprocessing using an identical bioinformatics pipeline, as described in “Methods”. Such practice not only mitigates batch effects and ensures consistency^17,18^, but also bolsters the identification of health- and disease-related gut taxonomic signatures despite the presence of potentially strong confounding factors. Indeed, this is supported by principal component analysis (PCA), where, despite the samples originating from varying sources and conditions, the healthy and non-healthy groups display significantly distinct gut microbiome profiles (R^2^ = 1.2%, *P* = 0.001, PERMANOVA; **Fig. 2a**).

**Figure 2.**
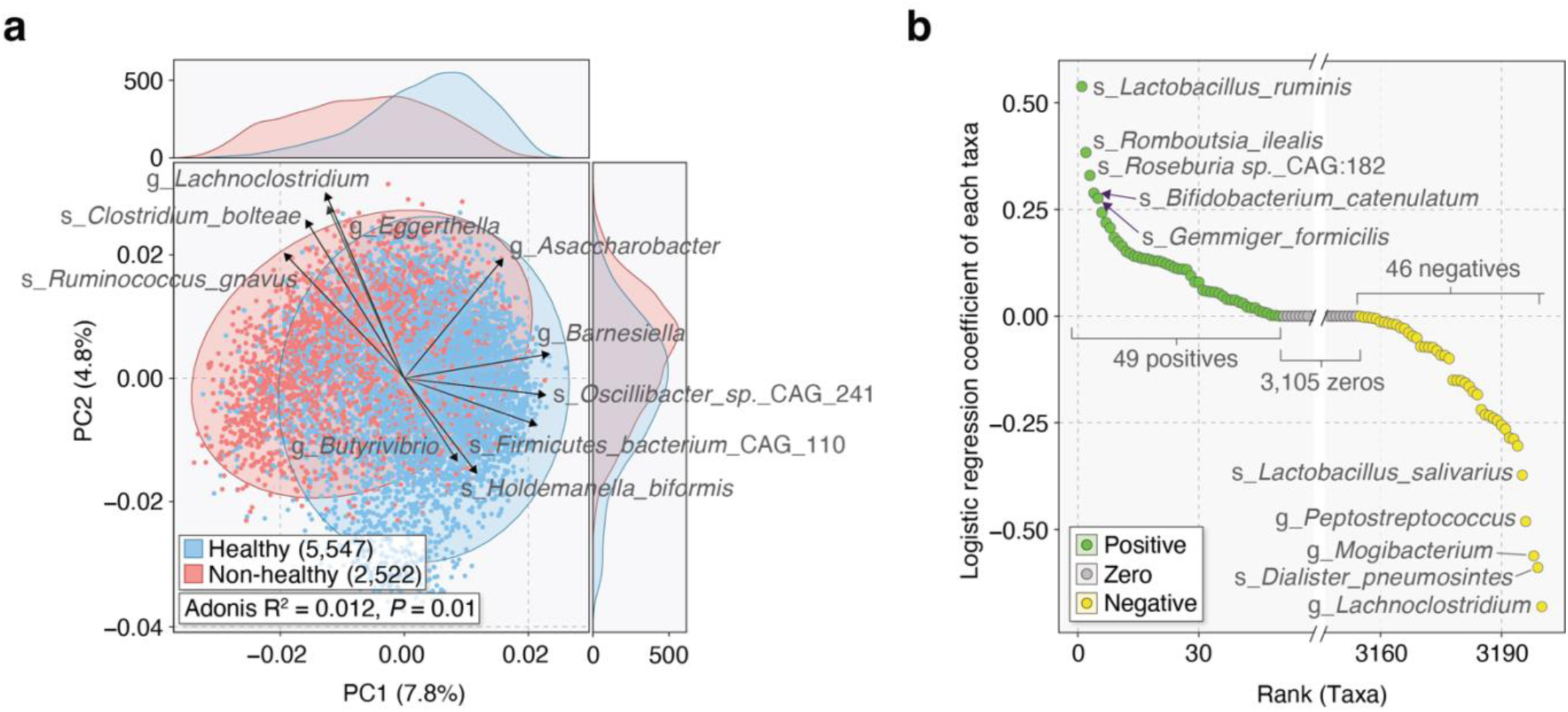
Gut microbiome taxonomic profiles of healthy and non-healthy individuals inform a Lasso-penalized logistic regression classification model. **(a)** Principal component analysis (PCA) of gut microbiome profiles reveals significant differences in the distribution of healthy (disease-free) (blue, n = 5547) and non-healthy (diseased) (red, n = 2522) groups (*P* < 0.05, PERMANOVA). Ellipses represent 95% confidence regions. The top 10 PC1 and PC2 loading vector magnitudes are shown. **(b)** Coefficient values for the Lasso-penalized logistic regression model. The model includes 49 taxa with positive coefficients, 3105 taxa with zero coefficients, and 46 taxa with negative coefficients.

### Implementing Lasso-penalized logistic regression in GMWI2

For the classification task, GMWI2 employs a Lasso-penalized logistic regression model instead of the log-ratio equation utilized in the original GMWI. Hence, GMWI2 essentially uses linear regression for its predictions, resembling polygenetic risk score models in statistical genetics^19,20^. The model was trained on gut microbiome taxonomic profiles (derived from the aforementioned pooled dataset of 8069 stool shotgun metagenomes) spanning all measurable taxonomic ranks to model disease likelihood as a linear function of microbial presence or absence. Specifically, the GMWI2 score for an individual sample is defined as the predicted log-odds (logit) of the sample originating from a healthy, non-diseased individual. A more comprehensive explanation of how GMWI2 employs Lasso-penalized logistic regression to estimate disease likelihood is detailed in “Methods”.

The original GMWI approach employed a prevalence-based strategy, identifying health and disease-associated microbial species based on differential occurrence frequencies. Our current method learns variable feature importances, obviating the need for manual species identification. More specifically, the Lasso-penalized logistic regression model identified 95 microbial taxa with non-zero coefficients (**Fig. 2b** and **Supplementary Data 3**). Interestingly, the majority of taxa characterized by positive and negative coefficients exhibited a higher relative abundance in the healthy and non-healthy groups, respectively (**Supplementary Data 4**). These identified taxa included 1 class, 3 orders, 4 families, 19 genera, and 68 species. Notably, the coefficient values varied between –0.68 to 0.54, ensuring that each taxon (i.e., clade) contributes differently to the GMWI2 score according to its relative association strength. This presents a shift from our previous GMWI log-ratio model where equal weight was assigned to each species.

### Enhanced classification of healthy and non-healthy gut microbiomes with GMWI2

GMWI2 scores were calculated for metagenomes by applying the learned coefficients in computing the predicted log-odds. A positive GMWI2 value classifies the sample as healthy, indicating disease absence; while a negative GMWI2 value classifies it as non-healthy, denoting disease presence. A GMWI2 of 0 implies an equal presence (or absence) of all taxa with both positive and negative coefficients, thereby classifying the sample as neither healthy nor non-healthy. When evaluated on the training dataset (8069 samples), GMWI2 demonstrated a balanced accuracy of 79.9% (healthy correct classification rate: 79.2%, non-healthy correct classification rate: 80.6%) and a Cliff’s Delta (*d*) effect size of 0.75, significantly surpassing the balanced accuracy and Cliff’s Delta reported by our original GMWI model (71.8%, *d* = 0.63) and traditional species-level α-diversity indices (Shannon Index, Simpson Index, and richness) (**Fig. 3a** and **Supplementary Data 5**). Our results indicate that GMWI2 differentiates between healthy and non-healthy groups much more effectively than GMWI, although both indices were strongly correlated (Pearson’s *r* = 0.81; **Supplementary Fig. 1**). Moreover, we found that the gut microbiomes of healthy individuals exhibit significantly higher GMWI2 scores compared to the 11 studied disease phenotypes (**Fig. 3b**).

**Figure 3.**
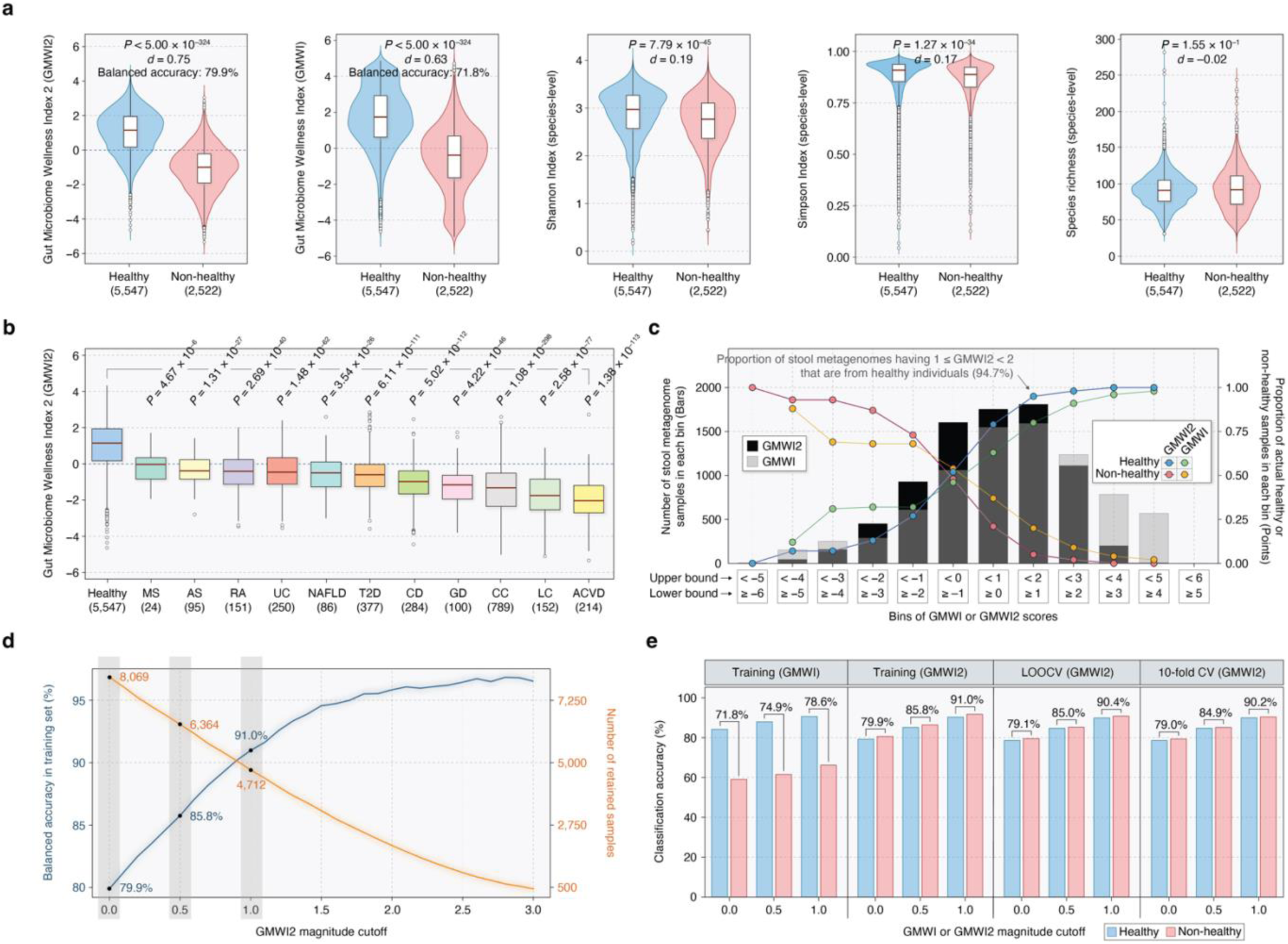
Enhanced classification of healthy and non-healthy stool metagenomes using Gut Microbiome Wellness Index 2 (GMWI2). **(a)** GMWI2 best stratifies healthy and non-healthy groups compared to GMWI and α-diversity indices (*d*, Cliff’s Delta effect size; *P*-values from the Mann-Whitney *U* test). Balanced accuracies on the training set are shown for GMWI2 and GMWI. **(b)** The healthy group (blue, far left) exhibits significantly higher GMWI2 scores than all 11 non-healthy phenotypes. **(b)** Bins of GMWI2 and GMWI scores (x-axis). The height of black and gray bars indicate metagenome sample counts in each GMWI2 and GMWI bin, respectively (y-axis, left). Points represent the proportion of samples in each GMWI2 or GMWI bin corresponding to actual healthy and non-healthy individuals (y-axis, right). **(d)** Increased magnitude cutoffs result in improved classification performance of GMWI2, showing increasing training set balanced accuracy (blue, y-axis, left) at the expense of decreasing retained samples (orange, y-axis, right). **(e)** Classification performances of GMWI and GMWI2 in distinguishing healthy and non-healthy groups. Balanced accuracies are depicted for both groups on the training set, leave-one-out cross-validation (CV), and 10-fold CV, using varying magnitude cutoffs (0, 0.5, 1.0) of GMWI and GMWI2 scores. Balanced accuracies are shown between the blue and pink bars, which represent healthy and non-healthy groups, respectively. For the 10-fold CV, repeated random sub-sampling was performed 10 times, and the average results are displayed.

We subsequently explored whether higher (or more positive) GMWI2 values could indicate enhanced confidence in categorizing stool metagenomes as healthy. Conversely, we examined if lower (or more negative) GMWI2 scores suggest an increased likelihood that a sample could be classified as non-healthy. Indeed, we observed a progressive increase in the proportion of healthy individuals among metagenome samples with increasingly positive GMWI2 scores. Similarly, increasingly negative GMWI2 scores captured larger proportions of the non-healthy subjects (**Fig. 3c** and **Supplementary Table 1**), potentially indicating enhanced prediction confidence for samples with highly positive or negative GMWI2 scores. Notably, the proportions of actual healthy and non-healthy samples within the positive and negative bins of GMWI2, respectively, were higher compared to the analogous GMWI bins (refer to points in **Fig. 3c**). This difference in sample distributions between the GMWI2 and GMWI bins underscores GMWI2’s improved capability to differentiate between healthy and non-healthy samples.

Remarkable improvement in classification performance was observed when considering increasing cutoffs for the *magnitude* (or absolute value) of GMWI2 scores, thereby signifying higher prediction confidence in the retained samples (**Supplementary Table 2**). For example, when retaining samples with GMWI2 magnitudes equal to or higher than 0.5 (i.e., GMWI2 scores below –0.5 or above +0.5) and 1.0 (i.e., GMWI2 scores below –1.0 or above +1.0), we achieved balanced accuracies of 85.8% and 91.0%, respectively (**Fig. 3d**). This approach, however, requires excluding samples with GMWI2 magnitudes below these cutoffs, leaving only 6364 (representing 78.9% of the total 8069 samples) and 4712 (58.4% of 8069) samples, respectively. These findings foster the potential utility of a “reject option”^21,22^ for low or zero GMWI2 magnitudes, which can serve as a criterion to redirect relatively uncertain predictions to other screening methods—this concept captures the understanding that certain aspects of health and disease are not fully explainable solely by the gut microbiome.

An important observation is that GMWI2 correctly classified healthy and non-healthy stool metagenomes at nearly the same rate (79.2% and 80.6%, respectively) despite imbalanced sample numbers. This contrasts markedly with the original GMWI, which achieved a much higher correct classification rate on healthy samples (**Fig. 3e**). We also assessed the performance of the GMWI2 model utilizing both leave-one-out cross-validation (LOOCV) and 10-fold cross-validation (10-fold CV) (**Fig. 3e**). Interestingly, GMWI2 achieved nearly identical balanced accuracies of 79.1% (healthy correct classification rate: 78.6%, non-healthy correct classification rate: 79.5%) and 79.0% (healthy correct classification rate: 78.6%, non-healthy correct classification rate: 79.3%) in LOOCV and 10-fold CV, respectively, nearly matching the performance achieved on the training dataset (79.9%). This outcome highlights the capability of the GMWI2 model to generalize effectively to new clinical settings regardless of the study (or site) specific characteristics.

To assess the impact of the aforementioned reject option, we computed classification performances using different magnitude cutoffs and cross-validation methods (**Fig. 3e**). Remarkably, GMWI2 achieved a balanced accuracy of 90.4% and 90.2% in LOOCV and 10-fold CV, respectively, on the samples with scores below –1.0 or above +1.0. These balanced accuracies were very close to those observed in the training set (91.0%). In contrast, when applying the same reject option criteria to GMWI (i.e., cutoff of 1.0), the balanced accuracy drops considerably to 78.6%. In all, these results emphasize the notable improvements achieved with GMWI2 over GMWI.

### Evaluating the robustness of GMWI2 across different study populations

Evaluation of any biomarker or molecular signature on independent samples is the gold standard for assessing its robustness^18^. We therefore conducted inter-study validation (ISV) to assess the impact of batch effects (i.e., technical or biological variations associated with study populations) on GMWI2 performance stability. In this approach, we iteratively excluded a single study, trained the GMWI2 model on the remaining studies, and evaluated its classification performance on the excluded study^23^. (The excluded study essentially becomes the independent cohort.) Despite the wide variability in classification performance across different studies (see gold points indicating ISV classification accuracy per study in **Fig. 4a** and **Supplementary Table 3**), the average balanced accuracy was 75.8%. This performance rose to 86.9% when considering samples with GMWI2 scores lower than –1 or higher than 1. The classification performances obtained from ISV exhibited minimal disparity compared to the performances achieved by LOOCV and 10-fold CV, which do not consider study boundaries; the small discrepancy between these strategies showcases GMWI2’s resilience against batch-related biases, indicating that GMWI2 generalizes effectively across stool metagenomes, regardless of the subjects’ origins. Further evidence of this robustness is demonstrated by the area-under-the-curve (AUC) metrics in the training set, 10-fold CV, and ISV, achieving AUCs of 0.88, 0.87, and 0.84, respectively (**Fig. 4b**).

**Figure 4.**
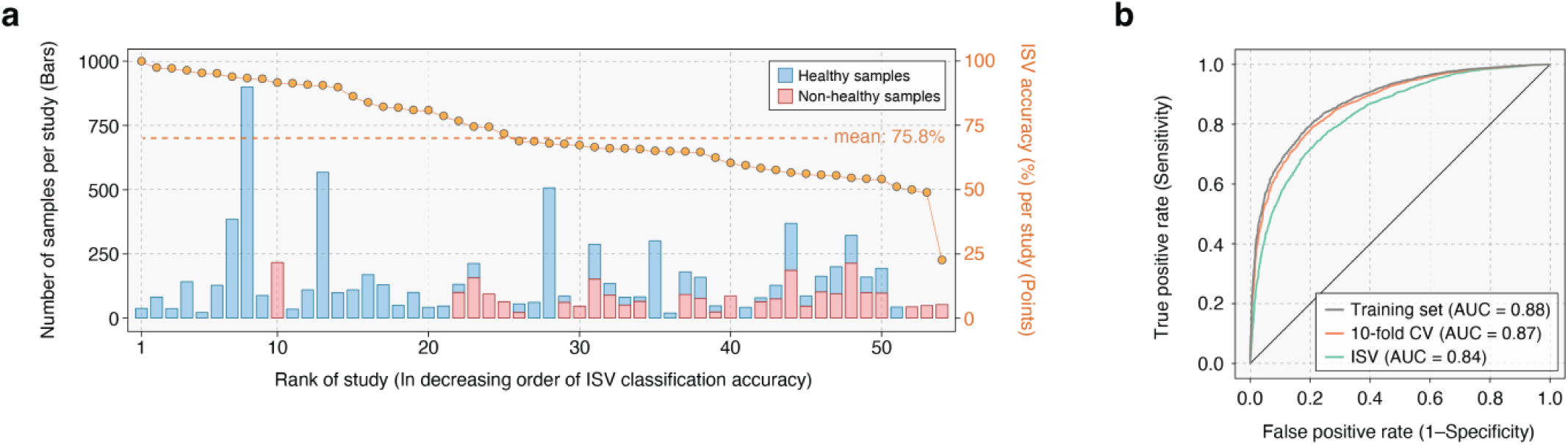
GMWI2 demonstrates effective generalization across diverse study populations. **(a)** Classification performance on each excluded study in inter-study validation (ISV) is displayed by gold points (y-axis, right). The studies on the x-axis are rank-ordered based on either classification performance for a single phenotype (healthy or non-healthy) or balanced accuracy in the case of both phenotypes. The stacked bars illustrate the number of healthy (blue) and non-healthy (pink) stool metagenome samples in each study (y-axis, left). **(b)** Receiver operating characteristic curves for classification performance in distinguishing healthy and non-healthy phenotypes on the training set, 10-fold CV, and ISV.

### Utilizing GMWI2 for new insights from existing longitudinal gut microbiome studies

Reanalyzing and reinterpreting previously studied datasets can be immensely valuable to gain new insights from existing data. Therefore, we reapplied GMWI2 to stool metagenomes from four recently published longitudinal gut microbiome studies to assess the model’s applicability across different datasets. Notably, these samples were not part of the initial pool of 8069 metagenomes used to train GMWI2. Using data from a case study^24^, we analyzed stool metagenomes from 22 individuals with irritable bowel syndrome (IBS) before and six months after receiving fecal microbiota transplantation (FMT) from two healthy donors. Among the participants, fourteen reported symptom relief after FMT (“Effect” group), while eight did not experience symptom relief (“No Effect” group) despite both groups demonstrating a significant increase in species richness at six months following the FMT (*P* < 0.05, one-tailed Wilcoxon signed-rank test; **Supplementary Fig. 2**). However, only the individuals in the “Effect” group exhibited a significant increase in GMWI2 (*P* < 0.05; **Fig. 5a** and **Supplementary Table 4**), indicating that GMWI2 may accurately capture the clinical response to treatment. Likewise, an increase in the species-level Shannon Index was observed only in the “Effect” group (*P* < 0.05; **Supplementary Fig. 3**). Overall, these findings suggest that while α-diversity metrics, such as richness and Shannon diversity, may yield conflicting conclusions, GMWI2 provides more direct relevance to subject phenotype following FMT treatment for IBS. Considering the ongoing concerns about the long-term safety and efficacy of FMT in treating patients with chronic diseases^25^, computational tools like GMWI2 could be useful in assisting with the selection of healthy donors and stool samples.

**Figure 5.**
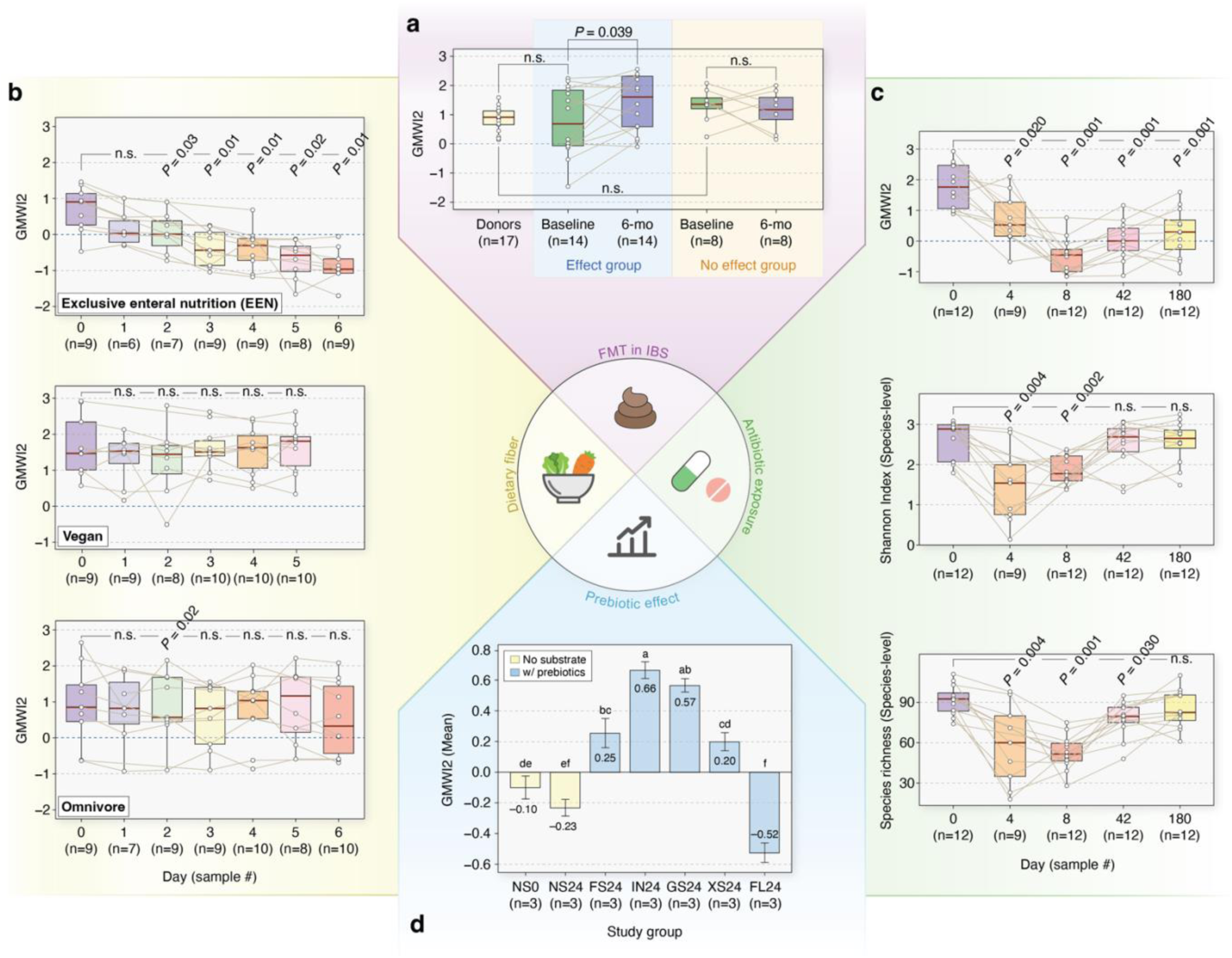
Reanalysis of existing longitudinal gut microbiome studies with GMWI2. **(a)** Changes in GMWI2 in patients with irritable bowel syndrome observed six months (6-mo) after undergoing fecal microbiota transplantation. Only subjects experiencing symptom relief (“Effect” group) displayed a significant increase in GMWI2 (*P* = 0.039, one-sided Wilcoxon signed-rank test). n, number of FMT donor samples (17 total samples from two healthy donors) or FMT recipients. **(b)** GMWI2 scores for the dietary groups (EEN, Vegan, and Omnivore) at baseline and the first 5 to 6 days of dietary intervention. The EEN group showed significant changes in GMWI2, with values significantly decreased by day 2 and thereafter (*P* < 0.05, two-sided Wilcoxon signed-rank test). No significant change in GMWI2 was observed for the Omnivore and Vegan groups compared to baseline. **(c)** GMWI2, Shannon Index, and species richness before and after antibiotic intervention. Despite recovery in Shannon Index and species richness at day 42 and 180, respectively, GMWI2 remained significantly lower compared to day 0, suggesting incomplete gut microbiome recovery even after ∼6 months (*P* < 0.05, two-sided Wilcoxon signed-rank test). **(d)** GMWI2 of microbial communities after 24-hour *in vitro* fecal fermentation of five different prebiotic oligosaccharides. The height of the bars represents the mean GMWI2 (experiment conducted in triplicates for each study group), and error bars indicate the standard deviation from the mean. Different small letters denote groups with significant differences in GMWI2 as determined by Tukey’s HSD test (*P* < 0.05). Control groups: NS0, no substrate addition at 0 h; NS24, no substrate for 24 h. Prebiotic groups: FS24, fructooligosaccharide; IN24, inulin; GS24, galactooligosaccharide; XS24, xylooligosaccharide; FL24, 2’-fucosyllactose.

In the second case study^26^, we investigated the effects of one’s diet on gut microbiome composition. We calculated GMWI2 for stool metagenomes obtained from 30 healthy volunteers before and during a dietary intervention. Three groups of participants were studied: Vegan (self-reported vegans who resumed their regular diet), Omnivore (participants who consumed a standard diet of both animal and plant origin), and Exclusive Enteral Nutrition (EEN) (participants with an omnivorous diet who consumed a synthetic, fiber-free diet for the duration of the study). Stool samples were collected at baseline and each day during the dietary intervention. We observed that the GMWI2 scores for both the vegan and omnivore subjects remained relatively stable throughout the intervention period of five to six days. However, GMWI2 for the EEN group significantly decreased relative to baseline by day 2 and onwards (*P* < 0.05, two-tailed Wilcoxon signed-rank test; **Fig. 5b** and **Supplementary Table 5**) while α-diversities did not significantly change across the groups (**Supplementary Fig. 4**). These results suggest that the removal of dietary fiber may lead to a rapid decrease in overall gut health, an early change detected solely by GMWI2 and not by α-diversity metrics. Overall, our findings strengthen the evidence for the well-established benefits of dietary fiber on health^27–29^ and suggest that changes in GMWI2 could effectively monitor these benefits over time.

For the third case study^30^, we calculated GMWI2 for stool metagenomes from twelve healthy young adults who underwent a 4-day exposure with broad-spectrum antibiotics (meropenem, gentamicin, and vancomycin). Here, stool samples were collected before the exposure, and then again at 4, 8, 42, and 180 days post-intervention. While species-level α-diversity measures (Shannon Index and richness) indicated that the gut microbiome may have recovered somewhat by day 42, GMWI2 did not demonstrate any recovery trend even by day 180 (**Fig. 5c** and **Supplementary Table 6**). These findings reflect deleterious post-intervention taxonomic shifts originally noted by Palleja *et al*., such as the rise in previously undetectable *Clostridium spp*., and the disappearance of probiotic members of *Bifidobacterium* and butyrate producers *Coprococcus eutactus* and *Eubacterium ventriosum*. Our results therefore offer a novel perspective on the long-term impact of short-term broad-spectrum antibiotic intervention on gut microbiota, and suggest that GMWI2 could be a valuable tool for clinicians in assessing gut microbiome recovery following an acute illness.

In the final case study^31^, we examined the effect of various oligosaccharides on gut microbial communities. In the original study, Lee *et al.* used GMWI as a quantitative tool to assess the prebiotic effect of oligosaccharides, with broader implications for designing personalized diets based on their impact on gut microbiome wellness. For the study, nineteen healthy adult volunteers (fourteen men and five women) provided fecal samples, which were then pooled. Fructooligosaccharides (FOS), galactooligosaccharides (GOS), xylooligosaccharides (XOS), inulin (IN), and 2’-fucosyllactose (2FL) were separately mixed with the fecal samples in a 24-hour *in vitro* anaerobic batch fecal fermentation system. Two control groups were also included: one without substrate addition at 0 hours (NS0) and another without substrate addition for 24 hours (NS24). The experiment was conducted in triplicates for each of the seven study groups.

GMWI2 was calculated for each of the 21 pooled fecal samples (**Fig. 5d** and **Supplementary Table 7**), thereby replicating the original study with our new index. Consistent with previous findings, the NS24 group exhibited a lower average GMWI2 than the NS0 group, indicating a less healthy, disease-associated state. Notably, the addition of the three prebiotics (FOS, IN, and GOS) resulted in significantly higher GMWI2 compared to NS0 (*P* < 0.05, Tukey’s HSD test). These same three prebiotics, along with XOS, led to significantly higher GMWI2 relative to NS24 (*P* < 0.05). However, unlike the GMWI2 results, traditional α-diversity metrics such as the Shannon Index, species richness, species evenness, and inverse Simpson’s Index showed significantly lower values in all prebiotic treatment groups compared to the NS0 group (*P* < 0.05). Therefore, at least in the *in vitro* fermentation setting, intake of these four prebiotics could potentially stimulate the growth of gut microbial species associated with healthy conditions, an effect observed solely by using GMWI2.

## Discussion

Recent research into the human gut microbiome has highlighted its potential to inform the development of innovative tools for predictive healthcare^32–37^. In this regard, we introduce GMWI2, a robust predictor of health status based on gut microbiome taxonomic profiles that displays significant technological advances compared to its prototype (GMWI). Our extensive multi-study pooled analysis of 8069 stool shotgun metagenomes encompasses a diverse range of demographics, amplifying the biologically relevant signal linking gut taxonomies to human health. Delivering a cross-validation balanced accuracy of approximately 90% for higher confidence samples (i.e., outside the “reject option”), GMWI2 establishes its strong reliability as a classifier that distinguishes between healthy and non-healthy phenotypes. Moreover, this study exemplifies the transformative potential of widespread data-sharing in empowering robust machine-learning applications that can effectively overcome batch effects and biases^23,38,39^. Furthermore, by revisiting and reinterpreting data from previously published datasets, GMWI2 can offer novel perspectives even for the established understanding of the impact of dietary influences, antibiotic exposure, and FMT on the gut microbiome. Lastly, this study highlights the importance of extensive data sharing in fostering robust machine learning applications, and in demonstrating resilience to batch effects and algorithmic bias^40^.

Our study, while providing insights into the predictive capabilities of the gut microbiome, has some limitations that need to be acknowledged. First, the model could benefit from the inclusion of more intricate microbiome features such as species growth rates, strain details, and functional potential. Incorporating these important factors may improve predictive accuracy and offer a richer perspective on the intricate mechanisms tying the gut microbiome to overall human health. Second, our dataset, while extensive, is limited in its diversity. By broadening our selection criteria, we could incorporate more varied metagenomes, capture a broader range of disease phenotypes (like neurodegenerative and psychiatric disorders), and reach a more diverse demographic. Such expansion could enhance the model’s generalizability across populations. Third, although we employed a species-level approach, there’s a potential missed opportunity in not focusing on microbial strains, which often bear more clinical significance. While our method surpasses the genus-level limitations of 16s rRNA gene amplicon sequencing, it doesn’t account for the variability among strains of the same species. Last, we made concerted efforts to ensure a diverse representation of geographies, ethnicities/races, and cultures. However, achieving a complete bias-free dataset is challenging. Future work should emphasize wider participant inclusion, especially from underrepresented areas and ethnicities, to truly globalize microbiome research.

GMWI2 is not for confirming a specific diagnosis of a disease, but rather to function as the proverbial “canary in a coal mine” by serving as an early warning system. It is designed to detect potentially adverse shifts in overall gut health, which could inform lifestyle modifications to prevent mild issues from escalating into severe health conditions, or prompt further diagnostic tests for a more detailed and accurate diagnosis. This could be particularly useful in clinical scenarios such as selecting FMT/organ donors, where gut health could be indicative of overall health, or in assessing early predictors of disease flare-ups. In conditions like rheumatoid arthritis and other autoimmune inflammatory disorders, GMWI2 could guide decisions on which patients might benefit from tapering or stopping therapy. In this sense, GMWI2 paves the way for a novel, transformative era in gut microbiome-centric health analytics, allowing for nuanced health evaluations tailored to individual microbial signatures. Looking ahead, integrating GMWI2 into a larger decision network alongside other biomeasurements (e.g., multi-omics, wearables) and AI-based models has the potential to open up exciting possibilities for healthy aging^41^ and preventative healthcare and wellness strategies^42,43^, driven by insights from our gut microbiome.

## Methods

### Multi-study pooling of human stool metagenomes

We conducted a comprehensive literature search using targeted keywords such as “gut microbiome”, “stool metagenome”, and “whole-genome shotgun” in PubMed and Google Scholar. The search was performed up until January 2022 to identify published studies that included publicly available shotgun metagenomic data of human stool samples, along with corresponding subject meta-data. In cases where multiple samples were collected from individuals across different time points, we included only the first or baseline sample from that study subject. Studies involving dietary or medication interventions, as well as those fewer than 40 samples, were not included in the pooled dataset for GMWI2 training. The raw sequence files (in .sra or .fastq format) were retrieved from the NCBI Sequence Read Archive and European Nucleotide Archive databases for further analysis.

### Stool metagenome sample exclusion criteria

To minimize potential bias and preserve data integrity, we applied stringent criteria to the stool shotgun metagenome samples included in our study. Specifically, we excluded samples sequenced using non-Illumina platforms, such as 454 GS FLX Titanium, Ion Torrent PGM, Ion Torrent Proton, and BGISEQ-500, to ensure consistency in sequencing technology. In terms of data quality, we excluded samples with low read counts (below 1 million reads) prior to quality control filtration. Additionally, our analysis did not include samples from studies with a primary focus on the virome or those where stool samples underwent virus-like particle purification.

Furthering our strict sample control standards, we also excluded disease control samples that were not specifically tied to a clinical diagnosis in the originating study. Individuals who were not specifically diagnosed with a specific disease but exhibited certain anomalous conditions were also excluded. These conditions comprised: (i) a Body Mass Index (BMI) suggestive of being underweight (BMI < 18.5), overweight (BMI ≥ 25 and < 30), or obese (BMI ≥ 30); (ii) declared heavy drug use (including alcohol and recreational drugs); (iii) age exceeding 100 years; and (iv) individuals initially healthy at baseline, but later reported to develop a disease condition during a longitudinal study. Lastly, we excluded non-healthy individuals with early-stage diseases (e.g., impaired glucose tolerance, hypertension, colorectal adenoma), rare or genetically-linked disorders (e.g., Behcet’s disease, schizophrenia), and non-colon cancers (including pancreatic, non-small cell lung, and breast cancer). These exclusions ensure a more uniform and representative dataset for training GMWI2, thereby enhancing the reliability and validity of our analyses.

### Quality control of sequenced reads

Potential human contamination was filtered out by removing reads that aligned to the human genome (reference genome GRCh38/hg38) using Bowtie2^44^ v2.4.4 with default parameters. Along with Illumina universal adapter sequences, probable adapter sequences were identified by extracting overrepresented sequences from each metagenome sample using FastQC^45^ v0.11.8. Adapter sequence clipping and quality filtration were performed using Trimmomatic^46^ v0.39. Specifically, Trimmomatic’s “ILLUMINACLIP” step was used, using a maximum seed mismatch count of 2, palindrome clip threshold of 30, simple clip threshold of 10, and minimum adapter length of 2 bp. Additionally, leading and trailing low-quality bases (Phred quality score < 3) of each read were removed, and trimmed reads shorter than 60 bp in nucleotide length were discarded.

### Taxonomic profiling

After performing quality filtration on all raw metagenomes, taxonomic profiling was carried out using the MetaPhlAn3^15^ v3.0.13 phylogenetic clade identification pipeline using default parameters. Briefly, MetaPhlAn3 classifies metagenomic reads to taxonomies based on a database (mpa_v30_CHOCOPhlAn_201901) of clade-specific marker genes. Once taxonomic features (or clades) of unknown/unclassified identity were removed, the remaining clades that could be detected in at least one metagenome sample in the pooled dataset were considered for further analysis.

After taxonomic profiling, the following metagenomes were discarded from our analysis: (i) samples composed of > 90% unmapped reads; (ii) samples with a relatively high proportion (> 25%) of unknown taxa; and (iii) samples lacking sufficient taxonomic diversity (< 100 identified taxa). These samples were removed to maintain quality and reliability of the training data. Finally, after applying all exclusion criteria, studies with fewer than 20 remaining samples were removed.

### Generating presence/absence taxonomic profiles

To mitigate concerns related to the compositional nature of microbiome data^47^, batch effects, and to simplify the interpretation of the GMWI2 classification model, we transformed the taxa relative abundances from MetaPhlAn3 into a binary presence/absence profile for each metagenome sample. Specifically, a taxon (i.e., clade) was deemed “present” in a given sample if its relative abundance in a sample was equal to or greater than 0.00001 (or 0.001%), and considered absent otherwise. Consequently, each sample was represented as a binary vector.

### PCA and PERMANOVA analysis on taxonomic profiles

Principal component analysis (PCA) was conducted on the presence/absence taxonomic profiles using the “prcomp” function in R. Furthermore, Bray-Curtis distance matrices were generated based on the relative abundances of microbial taxa (ranging from phylum to species) in the stool metagenomes. This was done using the “vegan” package v2.6.4 in R. We then carried out a permutational multivariate analysis of variance (PERMANOVA) on the distance matrix using the “adonis2” function. To evaluate the influence of the subjects’ health status on the total variance in gut microbial community composition, we calculated the *P*-value for the test statistic (pseudo-F) based on 999 permutations.

### Estimating disease likelihood using Lasso-penalized logistic regression

A Lasso-penalized logistic regression model (Python library “scikit-learn” v1.0.2) was trained on the binary presence/absence taxonomic profiles of the entire pooled dataset to predict disease presence. The L1 (Lasso) penalty was utilized with the LIBLINEAR solver^48^. The random state was set to 42, the regularization parameter *C* to 0.03, and class weight to “balanced”. The selection of the regularization parameter *C* was achieved through hyperparameter tuning, where we evaluated various candidates and selected the value (*C* = 0.03) that yielded the optimum classification performance in inter-study validation (ISV) (**Supplementary Table 8**). The class weight was set to “balanced” in order to account for the unbalanced class proportions in our pooled dataset.

Let ***x**_i_* be a binary vector encoding the presence or absence of *n* taxa in the *i^th^* labeled sample:

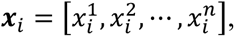

where 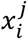 is 1 if taxa *j* is present in sample *i* and 0 otherwise. Additionally, *n* = 3200 is the number of taxonomic features (or clades) in the *i^th^* sample (a total of 3200 taxonomic features were observed at least once in the pooled metagenome dataset).

Let *y*_*i*_ represent the health status (1 for healthy, 0 for non-healthy) of sample *i*. The subsequent log-loss optimization objective function is solved using L1 regularization and class proportion weights as follows:

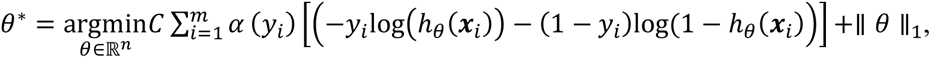

where *θ*^∗^ refers to the learned coefficient vector, *C* is the aforementioned inverse regularization strength parameter, *m* = 8069 represents the total number of samples in the pooled metagenome dataset, *⍺* is the class proportion weight term, and *h*_*θ*_(***x***_*i*_) is the hypothesis function:

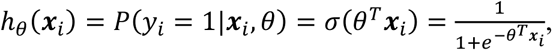

where *σ* is the sigmoid function. The class proportion term *⍺* accounts for the relatively unbalanced class proportions in the pooled dataset:

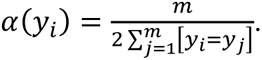

### Using GMWI2 as a stool metagenome-based health status classifier

We calculated GMWI2 scores for all 8069 stool metagenomes in the pooled dataset, as well as samples from the four gut microbiome case studies. The taxonomic profile of a metagenome was represented as a vector ***x***_*test*_, with binary values that encoded the presence or absence of microbial taxa. The computation employed the predicted log-odds (logit) using the previously learned coefficient vector *θ*^∗^:

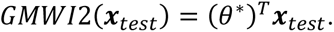

For classification purposes, a predetermined magnitude cutoff parameter *c* was utilized (*c* = 0 in case of having no cutoff or defer option). Finally, GMWI2 was computed on a metagenome ***x***_*test*_ while applying the following criteria:

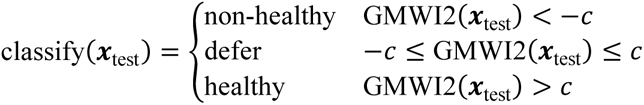

### Evaluation of classification performance

Balanced accuracy, defined as the average of the proportions of correctly classified healthy and non-healthy samples, was used to evaluate the performance of the GMWI2 classification model. This was done across different cutoff parameters (*c*) using multiple validation techniques: training on the entire dataset and then testing on the same training set, 10-fold cross-validation (10-fold CV), and leave-one-out cross-validation (LOOCV). In order to account for variability in 10-fold cross-validation, the process was repeated 10 times with shuffled fold partitions, and the results were averaged across all runs. Additionally, inter-study validation (ISV) was conducted, in which a single study was held out each time, the model was trained on the remaining studies, and testing was performed on the samples of the single held-out study. ISV allows for an assessment of classification performance across different studies.

## Data availability

All publicly available stool metagenome samples (and their corresponding studies) used in the analyses of this study are available in **Supplementary Data 2**. All raw metagenomic reads are available through SRA using the sequencing data accession IDs. Any additional information required to reanalyze the data reported in this paper is available from the corresponding author upon reasonable request.

## Code availability

A command-line tool for computing the GMWI2 score of a stool metagenome from its corresponding raw .fastq sequence file, the source code for the tool, processed datasets, and code used to generate figures and analyses, as well as complete instructions for installation and usage, will be freely available upon publication.

## Acknowledgements

We thank Professors Dan Knights and Konstantinos N. Lazaridis for helpful feedback regarding this study. This work was supported in part by the National Center for Advancing Translational Sciences of the National Institutes of Health Award Numbers UL1TR002494 and UL1TR002377. Additional support was provided by the Minnesota Partnership for Biotechnology and Medical Genomics through the Translational Product Development Fund (to J.S.), as well as Mark E. and Mary A. Davis to Mayo Clinic Center for Individualized Medicine (to J.S).

## Author contributions

D.C., V.K.G., and J.S. developed the study idea and designed all analytical methodologies. D.C. and V.K.G. performed the computational experiments. All authors analyzed and discussed the data. D.C., V.K.G., and J.S. wrote the manuscript, with contributions from other authors. All authors critically reviewed and approved the final manuscript.

