## Supplemental Data 1 for "Gut Microbiome Wellness Index 2 for Enhanced Health Status Prediction from Gut Microbiome Taxonomic Profiles"

for

Chang and Gupta *et al*.

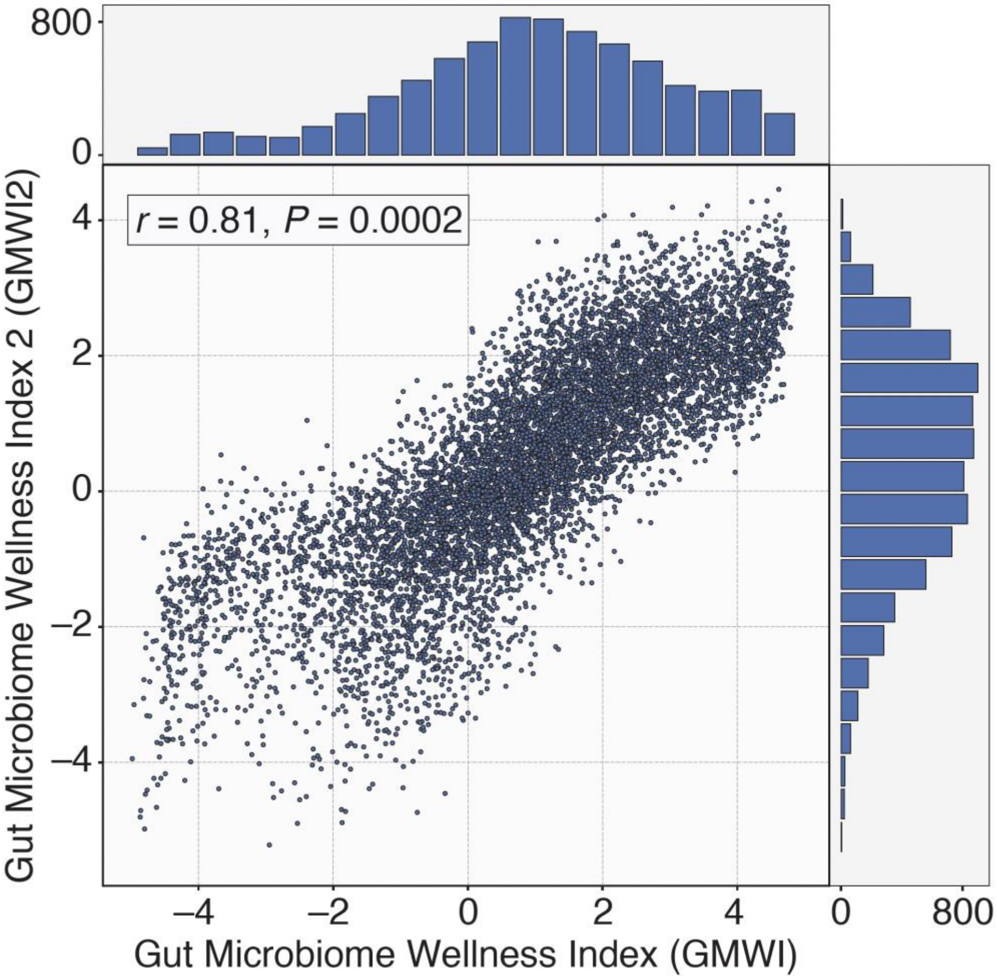

**Supplementary Figure 1. Correlation between GMWI and GMWI2 scores.** Scatter-plot showing the relationship between GMWI (x-axis) and GMWI2 (y-axis) calculated for each stool shotgun metagenome sample ($n$ = 8069). A strong correlation (Pearson's $r$ = 0.81, $P$ = 0.0002) between both versions of GMWIs was observed. Horizontal and vertical bars represent the number of scatter-plot points (i.e., samples) in each bin.

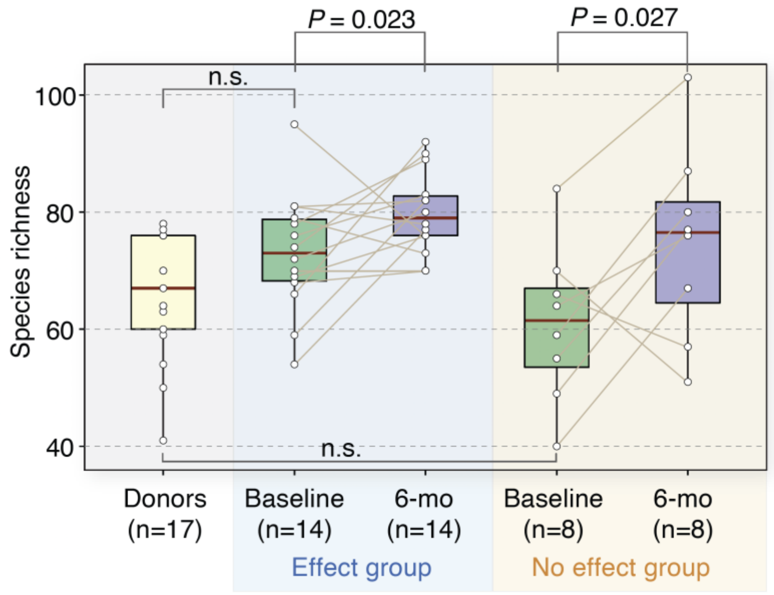

**Supplementary Figure 2. Species richness in gut microbiomes of patients with irritable bowel syndrome (IBS) observed before (Baseline) and six months after (6-mo) receiving fecal microbiota transplantation (FMT) therapy.** According to the original study (Goll, R. *et al.* Effects of Fecal Microbiota Transplantation in Subjects with Irritable Bowel Syndrome are Mirrored by Changes in Gut Microbiome. *Gut Microbes* 12, 1794263 (2020)), the "Effect group" subjects were those who experienced relief in IBS symptoms, whereas the "No effect group" subjects were those who did not. Significant increase in species richness was observed in both clinical groups ($P < 0.05$). $P$-values were calculated using the one-sided Wilcoxon signed-rank test. n.s., non-significant ($P \geq 0.05$). n, number of FMT donor samples or FMT recipients.

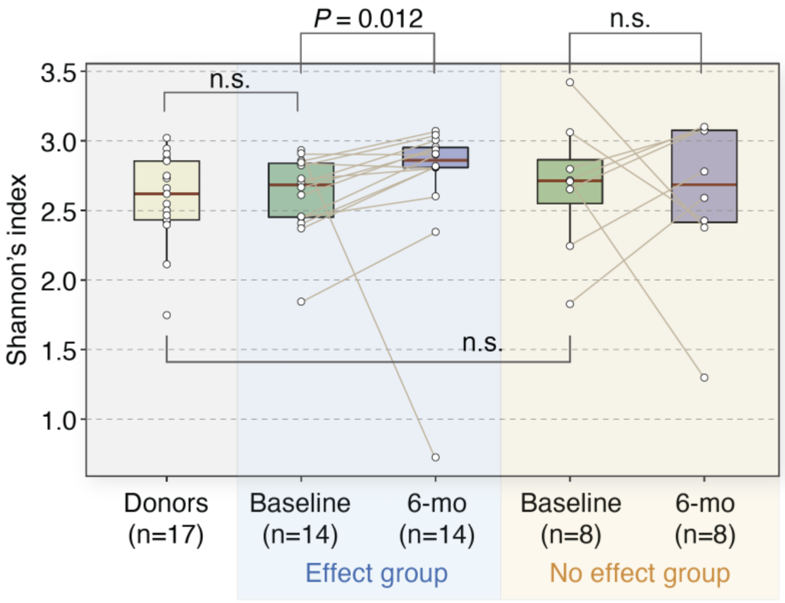

**Supplementary Figure 3. Shannon's Index in gut microbiomes of patients with irritable bowel syndrome (IBS) observed before (Baseline) and six months after (6-mo) receiving fecal microbiota transplantation (FMT) therapy.** According to the original study (Goll, R. *et al.* Effects of Fecal Microbiota Transplantation in Subjects with Irritable Bowel Syndrome are Mirrored by Changes in Gut Microbiome. *Gut Microbes* 12, 1794263 (2020)), the "Effect group" subjects were those who experienced relief in IBS symptoms, whereas the "No effect group" subjects were those who did not. Significant increase in Shannon's Index was observed in the "Effect group" only ($P < 0.05$). $P$-values were calculated using the one-sided Wilcoxon signed-rank test. n.s., non-significant ($P \geq 0.05$). n, number of FMT donor samples or FMT recipients.

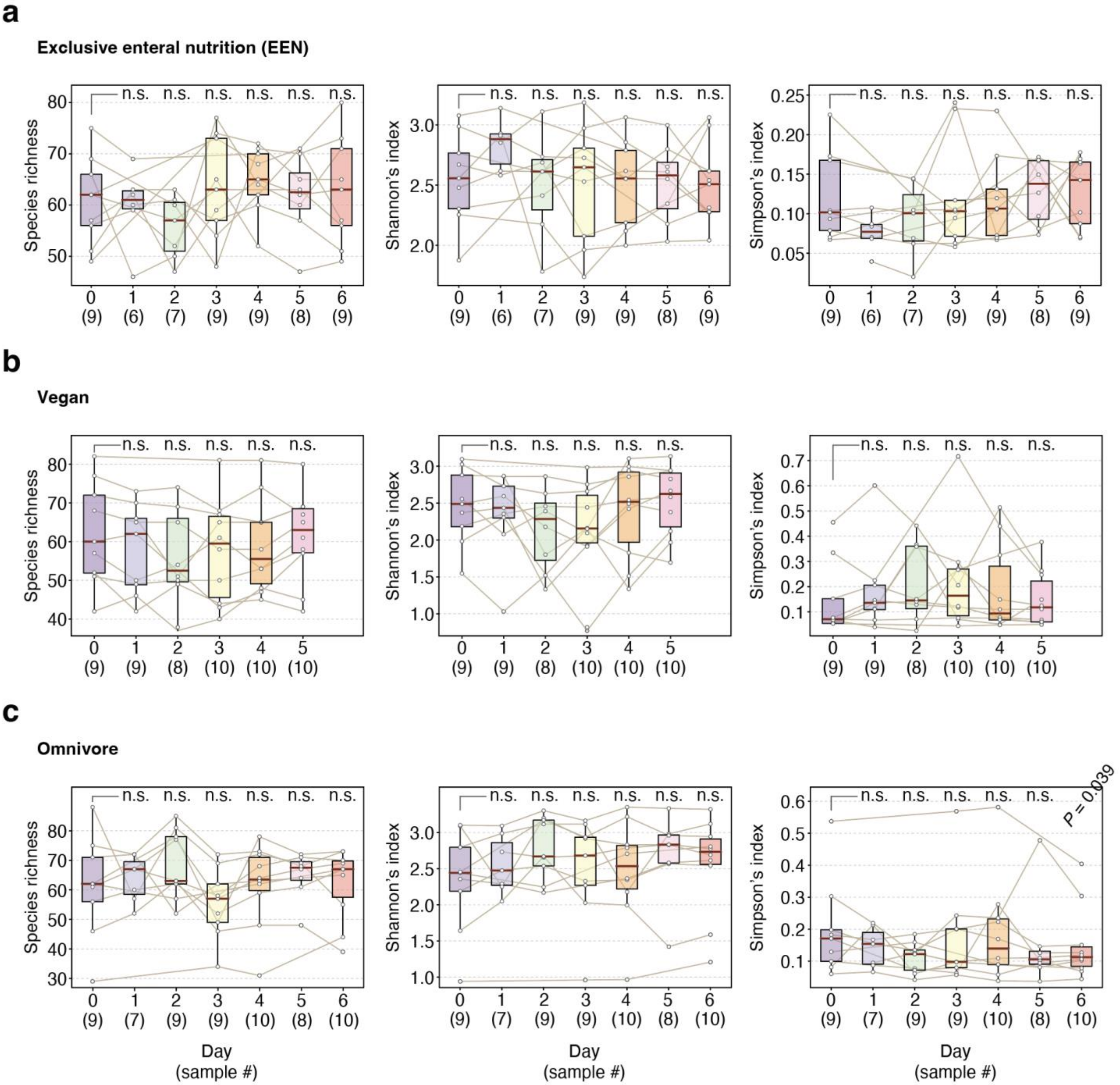

**Supplementary Figure 4. α-diversity indices across three diet groups.** Species richness, Shannon's Index, and Simpson's Index for the **(a)** Exclusive enteral nutrition (EEN) group, the **(b)** Vegan group, and the **(c)** Omnivore group at baseline and/or across the first five to six days of dietary intervention. For all diet groups, which are described in detail in the original study (Tanes, C. *et al.* Role of Dietary Fiber in the Recovery of the Human Gut Microbiome and its Metabolome. *Cell Host Microbe* 29, 394-407.e5 (2021)), α-diversity indices did not significantly change ($P \geq 0.05$; two-tailed Wilcoxon signed-rank test) compared to baseline (Day 0) except for Simpson's Index at Day 6 in the Omnivore group ($P = 0.039$).

**Supplementary Table 1.** Numbers (and proportions) of healthy and non-healthy samples in GMWI2 and GMWI bins. The table accounts for a total of 8,069 stool shotgun metagenome samples.

| Bins | GMWI2 | | GMWI | |
|---|---|---|---|---|
| | **Number of healthy samples (Proportion of total in bin)** | **Number of non-healthy samples (Proportion of total in bin)** | **Number of healthy samples (Proportion of total in bin)** | **Number of non-healthy samples (Proportion of total in bin)** |
| (–6, –5] | 0 (0.0%) | 4 (100.0%) | 0 (–) | 0 (–) |
| (–5, –4] | 3 (6.8%) | 41 (93.2%) | 19 (12.4%) | 134 (87.6%) |
| (–4, –3] | 11 (6.9%) | 148 (93.1%) | 78 (31.1%) | 173 (68.9%) |
| (–3, –2] | 57 (12.6%) | 394 (87.4%) | 91 (31.8%) | 195 (68.2%) |
| (–2, –1] | 252 (27.2%) | 676 (72.8%) | 195 (32.1%) | 413 (67.9%) |
| (–1, 0] | 833 (52.0%) | 770 (48.0%) | 484 (45.7%) | 575 (54.3%) |
| (0, 1] | 1380 (78.7%) | 374 (21.3%) | 975 (63.1%) | 571 (36.9%) |
| (1, 2] | 1713 (94.7%) | 96 (5.3%) | 1268 (80.4%) | 309 (19.6%) |
| (2, 3] | 1094 (98.4%) | 18 (1.6%) | 1123 (90.8%) | 114 (9.2%) |
| (3, 4] | 199 (99.5%) | 1 (0.5%) | 756 (96.4%) | 28 (3.6%) |
| (4, 5] | 5 (100.0%) | 0 (0.0%) | 558 (98.2%) | 10 (1.8%) |

**Supplementary Table 2.** Classification performance on training dataset for varying GMWI2 score magnitude cutoffs.

| GMWI2 cutoff (Magnitude) | Number of samples retained | Balanced accuracy[†] (%) |
|---|---|---|
| 0.0 | 8069 | 79.89 |
| 0.1 | 7719 | 81.19 |
| 0.2 | 7333 | 82.49 |
| 0.3 | 7004 | 83.53 |
| 0.4 | 6691 | 84.65 |
| 0.5 | 6364 | 85.75 |
| 0.6 | 6029 | 87.05 |
| 0.7 | 5664 | 88.20 |
| 0.8 | 5365 | 89.18 |
| 0.9 | 5049 | 90.17 |
| 1.0 | 4712 | 90.98 |
| 1.1 | 4412 | 91.63 |
| 1.2 | 4087 | 92.59 |
| 1.3 | 3782 | 93.35 |
| 1.4 | 3521 | 93.89 |
| 1.5 | 3238 | 94.56 |
| 1.6 | 2943 | 94.72 |
| 1.7 | 2697 | 95.00 |
| 1.8 | 2451 | 95.51 |
| 1.9 | 2210 | 95.53 |
| 2.0 | 1975 | 95.84 |
| 2.1 | 1756 | 96.07 |
| 2.2 | 1545 | 95.97 |
| 2.3 | 1347 | 96.10 |
| 2.4 | 1168 | 96.21 |
| 2.5 | 998 | 96.45 |
| 2.6 | 843 | 96.73 |
| 2.7 | 719 | 96.50 |
| 2.8 | 589 | 96.84 |
| 2.9 | 507 | 96.81 |
| 3.0 | 412 | 96.53 |

[†]Balanced accuracy is defined as the average of the proportions of healthy and non-healthy samples that were correctly classified.

**Supplementary Table 3.** GMWI2 classification performance on held-out studies in inter-study validation.

| Author's last name (Publication year) | Number of healthy samples | Number of non-healthy samples | Balanced accuracy[†] (%) |
|---|---|---|---|
| Smits (2017) | 38 | 0 | 100 |
| Jacobson (2021) | 82 | 0 | 97.56 |
| Lokmer (2019) | 37 | 0 | 97.30 |
| Pasolli (2019) | 142 | 0 | 96.48 |
| Ang (2021) | 22 | 0 | 95.45 |
| D'Souza (2021) | 128 | 0 | 95.31 |
| Schirmer (2016) | 385 | 0 | 94.03 |
| Zeevi (2015) | 900 | 0 | 93.44 |
| Le Chatelier (2013) | 88 | 0 | 93.18 |
| Yachida (2019) | 0 | 217 | 91.71 |
| Rettedal (2021) | 35 | 0 | 91.43 |
| Tett (2019) | 110 | 0 | 90.91 |
| Asnicar (2021) | 568 | 0 | 90.67 |
| De Filippis (2019) | 99 | 0 | 89.90 |
| Liu (2016) | 110 | 0 | 86.36 |
| Costea (2017) | 169 | 0 | 84.02 |
| Xie (2016) | 130 | 0 | 82.31 |
| Roager (2019) | 50 | 0 | 82.00 |
| Backhed (2015) | 100 | 0 | 81.00 |
| Yassour (2018) | 42 | 0 | 80.95 |
| Dhakan (2019) | 47 | 0 | 78.72 |
| Zhu (2021) | 32 | 100 | 76.81 |
| Franzosa (2018) | 56 | 157 | 74.57 |
| Gu (2017) | 0 | 94 | 74.47 |
| Ananthakrishnan (2017) | 0 | 64 | 71.88 |
| Wirbel (2019) | 33 | 22 | 68.94 |
| Kim (2021) | 61 | 0 | 68.85 |
| Huttenhower (2012) and Lloyd-Price (2017)[a] | 507 | 0 | 68.05 |
| Lloyd-Price (2019) | 25 | 61 | 67.87 |
| Feng (2015) | 0 | 46 | 67.39 |
| Qin (2014) | 135 | 152 | 66.59 |
| Zeller (2014) | 45 | 90 | 66.11 |
| Vogtmann (2016) | 30 | 51 | 66.08 |
| Schirmer (2018) | 18 | 65 | 65.81 |
| Mehta (2018) | 301 | 0 | 65.12 |
| Obregon-Tito (2015) | 20 | 0 | 65.00 |
| Yang (2020) | 88 | 92 | 64.90 |
| Nielsen (2014) | 82 | 77 | 64.68 |

| | | | |
|---|---|---|---|
| Ventura (2019) | 24 | 24 | 62.50 |
| Loomba (2017) | 0 | 86 | 60.47 |
| Sun (2021) | 42 | 0 | 59.52 |
| Weng (2019) | 15 | 64 | 58.44 |
| Yu (2015) | 53 | 75 | 57.71 |
| Qin (2012) | 183 | 186 | 56.68 |
| He (2017) | 40 | 46 | 56.25 |
| Zhang (2015) | 61 | 102 | 55.79 |
| Wen (2017) | 105 | 95 | 55.56 |
| Jie (2017) | 108 | 214 | 54.61 |
| Thomas (2019) | 61 | 99 | 54.27 |
| Yang (2021) | 97 | 97 | 54.12 |
| Qi (2019) | 43 | 0 | 51.16 |
| Davies (2020) | 0 | 44 | 50.00 |
| Gupta (2020) | 0 | 49 | 48.98 |
| Karlsson (2013) | 0 | 53 | 22.64 |

[a]Samples combined from both phases of the Human Microbiome Project (HMP1 and HMP1-II). [†]Balanced accuracy is defined as the average of the proportions of healthy and non-healthy samples that were correctly classified.

**Supplementary Table 4.** GMWI2 scores, species richness, and Shannon Index of stool metagenomes from Goll *et al*.

| Subject ID | Time-point[a] | GMWI2 | Species richness | Shannon Index | Study group[†] |
|---|---|---|---|---|---|
| 3 | Baseline | 1.97113 | 70 | 2.72843 | Effect |
| 3 | 6-mo | 1.91417 | 70 | 2.80772 | Effect |
| 11 | Baseline | –0.08548 | 81 | 2.45591 | Effect |
| 11 | 6-mo | 0.59796 | 78 | 2.60174 | Effect |
| 16 | Baseline | 0.01185 | 59 | 2.37104 | Effect |
| 16 | 6-mo | –0.10259 | 82 | 2.81796 | Effect |
| 19 | Baseline | 1.72307 | 54 | 2.40814 | Effect |
| 19 | 6-mo | 0.17037 | 77 | 2.81474 | Effect |
| 24 | Baseline | 0.15272 | 76 | 2.84407 | Effect |
| 24 | 6-mo | 2.39152 | 83 | 2.80925 | Effect |
| 25 | Baseline | –0.01581 | 66 | 2.91005 | Effect |
| 25 | 6-mo | 2.20480 | 92 | 2.90612 | Effect |
| 30 | Baseline | 1.87864 | 81 | 2.82448 | Effect |
| 30 | 6-mo | 2.55779 | 82 | 3.04217 | Effect |
| 31 | Baseline | 2.15370 | 74 | 2.85770 | Effect |
| 31 | 6-mo | 1.85034 | 90 | 3.07087 | Effect |
| 59 | Baseline | –0.13333 | 79 | 2.66701 | Effect |
| 59 | 6-mo | 1.35974 | 73 | 2.93682 | Effect |
| 77 | Baseline | –1.45895 | 95 | 2.93271 | Effect |
| 77 | 6-mo | 0.10312 | 76 | 0.72586 | Effect |
| 78 | Baseline | 1.48054 | 72 | 1.84437 | Effect |
| 78 | 6-mo | 1.03121 | 80 | 2.34608 | Effect |
| 88 | Baseline | 2.24283 | 69 | 2.70017 | Effect |
| 88 | 6-mo | 2.35412 | 76 | 3.00670 | Effect |
| 89 | Baseline | –0.53592 | 78 | 2.45049 | Effect |
| 89 | 6-mo | 0.58335 | 89 | 2.90201 | Effect |
| 90 | Baseline | 1.21949 | 68 | 2.61136 | Effect |
| 90 | 6-mo | 2.36111 | 70 | 2.95703 | Effect |
| 12 | Baseline | 1.47657 | 84 | 3.42045 | No effect |
| 12 | 6-mo | 0.14123 | 103 | 2.37705 | No effect |
| 18 | Baseline | 0.22944 | 70 | 2.70798 | No effect |
| 18 | 6-mo | 1.00296 | 51 | 1.29897 | No effect |
| 22 | Baseline | 1.31860 | 49 | 1.82744 | No effect |
| 22 | 6-mo | 1.98558 | 77 | 2.58996 | No effect |
| 32 | Baseline | 1.31242 | 40 | 2.24464 | No effect |
| 32 | 6-mo | 1.80714 | 67 | 2.77963 | No effect |
| 56 | Baseline | 1.36916 | 64 | 2.71700 | No effect |
| 56 | 6-mo | 0.27024 | 76 | 3.10007 | No effect |

| | | | | | |
|---|---|---|---|---|---|
| 64 | Baseline | 1.82462 | 55 | 2.79772 | No effect |
| 64 | 6-mo | 1.00534 | 80 | 3.07260 | No effect |
| 66 | Baseline | 0.83050 | 59 | 2.65193 | No effect |
| 66 | 6-mo | 1.50540 | 87 | 3.08256 | No effect |
| 75 | Baseline | 2.06776 | 66 | 3.06168 | No effect |
| 75 | 6-mo | 1.31720 | 57 | 2.42652 | No effect |
| D-1_S9 | - | 1.12769 | 76 | 2.85490 | Healthy donor |
| D-10_S4 | - | 0.69000 | 78 | 2.90576 | Healthy donor |
| D-12_S5 | - | 0.92771 | 70 | 2.61911 | Healthy donor |
| D-13U_S1 | - | 0.79794 | 41 | 2.65966 | Healthy donor |
| D-14_S8 | - | 0.18140 | 76 | 2.46545 | Healthy donor |
| D-14U_S10 | - | 0.14199 | 60 | 2.11311 | Healthy donor |
| D-15Fryst_S3 | - | 0.44763 | 59 | 2.87136 | Healthy donor |
| D-15U_S6 | - | 1.58681 | 67 | 2.49936 | Healthy donor |
| D-2_S7 | - | 1.08010 | 78 | 2.54714 | Healthy donor |
| D-3_S6 | - | 1.22138 | 77 | 2.94429 | Healthy donor |
| D-4_S10 | - | 1.14330 | 76 | 2.39656 | Healthy donor |
| D-5Fryst_S8 | - | 0.90896 | 64 | 2.43220 | Healthy donor |
| D-6Fresk_S5 | - | 0.86373 | 64 | 3.02126 | Healthy donor |
| D-6Fryst_S4 | - | 1.10229 | 54 | 2.74909 | Healthy donor |
| D-7Fryst_S7 | - | 1.35535 | 70 | 2.42049 | Healthy donor |
| D-9Feresk_S9 | - | 0.24005 | 50 | 1.74761 | Healthy donor |
| D-9Fryst_S2 | - | 0.65847 | 63 | 2.73193 | Healthy donor |

[a]Before (Baseline) and six months after (6-mo) receiving fecal microbiota transplantation (FMT) from Healthy donors (-).
[†]According to the original study, “Effect group” subjects were those who experienced relief in IBS symptoms, whereas the “No effect group” subjects were those who did not.

**Supplementary Table 5.** GMWI2 scores of stool metagenomes from Tanes *et al*.

| Diet group | Subject ID | Time-point[†] | | | | | | |
|---|---|---|---|---|---|---|---|---|
| | | Day 0 | Day 1 | Day 2 | Day 3 | Day 4 | Day 5 | Day 6 |
| EEN[a] | 9003 | 0.934 | 0.483 | 0.501 | –0.58 | –0.59 | - | –0.962 |
| | 9009 | 0.905 | - | - | –0.437 | –0.722 | –0.689 | –0.863 |
| | 9013 | 1.388 | - | 0.256 | 0.253 | –0.041 | –1.66 | –1.077 |
| | 9016 | 1.141 | - | 0.903 | 0.06 | - | –0.466 | –0.679 |
| | 9024 | –0.472 | –0.28 | –0.577 | –1.043 | –1.179 | –1.24 | –1.701 |
| | 9025 | 1.461 | 1.008 | - | - | 0.682 | - | - |
| | 9029 | 0.264 | 0.075 | 0.009 | –0.101 | –0.308 | –0.389 | –0.339 |
| | 9032 | 0.525 | - | - | 0.228 | –0.146 | –0.142 | –0.055 |
| | 9034 | 0.14 | –0.014 | –0.05 | –0.862 | –0.109 | –0.125 | –1.077 |
| | 9038 | - | –0.308 | –0.728 | –0.917 | –1.118 | –0.961 | –1 |
| Omnivore | 9005 | –0.624 | - | - | –0.897 | –0.624 | - | –0.607 |
| | 9006 | 2.645 | 0.818 | 2.153 | 1.436 | 2.022 | 1.651 | 1.14 |
| | 9010 | 0.849 | 0.647 | 0.57 | –0.175 | 0.545 | 0.269 | 0.599 |
| | 9015 | 0.668 | 0.118 | 0.367 | 0.816 | 1.029 | –0.2 | –0.691 |
| | 9019 | 1.472 | 1.915 | 1.693 | 1.402 | 1.73 | 1.686 | 1.533 |
| | 9030 | 1.276 | - | 0.513 | 0.539 | 1.275 | 0.677 | 0.048 |
| | 9033 | –0.636 | -0.929 | –0.899 | –0.362 | –0.864 | –0.589 | –0.589 |
| | 9036 | 0.452 | - | 0.444 | - | 0.526 | - | 0.029 |
| | 9037 | 2.205 | 1.86 | 1.68 | 1.552 | 1.044 | 2.212 | 2.082 |
| | 9040 | - | 1.225 | 1.4 | 1.331 | 1.295 | 1.708 | 1.662 |
| Vegan | 9002 | 0.57 | 0.396 | 0.423 | 0.921 | 0.658 | 0.337 | - |
| | 9004 | 0.914 | 1.492 | 1.345 | 1.393 | 1.619 | 0.96 | - |
| | 9008 | 1.602 | 1.451 | 1.77 | 1.549 | 1.77 | 1.605 | - |
| | 9014 | 1.468 | 1.569 | –0.509 | 1.405 | 0.884 | 1.849 | - |
| | 9017 | 1.008 | 0.161 | 1.054 | 0.726 | 0.498 | 0.89 | - |
| | 9022 | - | 1.742 | 1.537 | 1.887 | 2.018 | 1.943 | - |
| | 9026 | 2.342 | 2.128 | - | 1.462 | 1.634 | 1.76 | - |
| | 9027 | 2.92 | - | - | 2.486 | 2.433 | 2.627 | - |
| | 9031 | 1.458 | 1.756 | 1.663 | 1.609 | 1.574 | 1.886 | - |
| | 9035 | 2.892 | 3.045 | 2.796 | 2.621 | 2.371 | 2.277 | - |

[a]Exclusive Enteral Nutrition (EEN), which is a synthetic diet devoid of fiber. [†]Baseline and/or across the first five to six days of dietary intervention. '-' denotes a time-point where a stool metagenome sample was unavailable.

**Supplementary Table 6.** GMWI2 scores, species richness, and Shannon Index of stool metagenomes from Palleja *et al.*

| Index | Subject ID | Time-point† | | | | |
|---|---|---|---|---|---|---|
| | | Day 0 | Day 4 | Day 8 | Day 42 | Day 180 |
| GMWI2 | ERAS1 | 2.589 | - | –0.506 | –0.022 | 0.058 |
| | ERAS2 | 1.083 | 0.145 | –0.994 | –0.123 | –0.577 |
| | ERAS3 | 0.889 | 0.526 | –1.141 | –0.45 | –0.663 |
| | ERAS4 | 1.442 | 2.103 | –1.152 | –1.126 | –0.167 |
| | ERAS5 | 0.98 | –0.67 | –0.399 | 0.525 | 0.53 |
| | ERAS6 | 2.929 | 0.758 | 0.765 | 0.678 | 1.598 |
| | ERAS7 | 1.936 | 0.161 | –0.87 | –0.662 | 0.301 |
| | ERAS8 | 0.985 | - | 0.184 | 1.164 | 0.567 |
| | ERAS9 | 2.101 | 1.734 | –0.346 | 0.38 | 1.048 |
| | ERAS10 | 2.518 | - | –1.006 | –0.258 | –1.046 |
| | ERAS11 | 2.455 | 1.269 | –0.33 | 0.027 | 1.197 |
| | ERAS12 | 1.589 | 0.344 | –0.075 | 0.218 | 0.288 |
| Species richness | ERAS1 | 74 | - | 59 | 76 | 81 |
| | ERAS2 | 89 | 18 | 53 | 72 | 78 |
| | ERAS3 | 103 | 96 | 28 | 48 | 82 |
| | ERAS4 | 111 | 98 | 75 | 81 | 110 |
| | ERAS5 | 95 | 60 | 47 | 77 | 72 |
| | ERAS6 | 82 | 23 | 39 | 87 | 97 |
| | ERAS7 | 93 | 80 | 70 | 80 | 83 |
| | ERAS8 | 84 | - | 61 | 90 | 95 |
| | ERAS9 | 92 | 71 | 50 | 86 | 95 |
| | ERAS10 | 93 | - | 49 | 59 | 70 |
| | ERAS11 | 106 | 46 | 45 | 95 | 99 |
| | ERAS12 | 78 | 35 | 57 | 79 | 61 |
| Shannon Index | ERAS1 | 2.886 | - | 1.644 | 2.785 | 2.534 |
| | ERAS2 | 3.035 | 0.77 | 1.622 | 2.619 | 2.562 |
| | ERAS3 | 3.035 | 1.636 | 1.428 | 2.223 | 2.545 |
| | ERAS4 | 2.996 | 2.897 | 2.338 | 2.895 | 2.995 |
| | ERAS5 | 2.083 | 0.915 | 1.584 | 2.543 | 2.77 |
| | ERAS6 | 2.013 | 0.152 | 1.394 | 3.066 | 3.264 |
| | ERAS7 | 2.909 | 1.553 | 2.389 | 2.372 | 1.504 |
| | ERAS8 | 2.075 | - | 2.091 | 1.337 | 2.079 |
| | ERAS9 | 2.934 | 2.812 | 1.745 | 2.952 | 2.779 |
| | ERAS10 | 2.668 | - | 1.834 | 1.47 | 1.813 |
| | ERAS11 | 3.101 | 2.012 | 2.239 | 2.822 | 3.139 |
| | ERAS12 | 1.806 | 0.651 | 2.224 | 3.078 | 2.826 |

†Before (0) and at 4, 8, 42, and 180 days post-intervention with a cocktail of three last-resort antibiotics (meropenem, gentamicin, and vancomycin). '-' denotes a time-point where a stool metagenome sample was unavailable.

**Supplementary Table 7.** GMWI2 scores of pooled[†] fecal samples from Lee *et al*.

| Study groups[a] | Replicate 1 | Replicate 2 | Replicate 3 |
|---|---|---|---|
| NS0 | –0.117 | –0.22 | 0.038 |
| NS24 | –0.177 | –0.177 | –0.340 |
| FS24 | 0.065 | 0.350 | 0.350 |
| IN24 | 0.725 | 0.556 | 0.713 |
| GS24 | 0.611 | 0.608 | 0.478 |
| XS24 | 0.192 | 0.303 | 0.100 |
| FL24 | –0.607 | –0.400 | –0.568 |

[†]Nineteen healthy adult volunteers (14 males and 5 females) provided fecal samples, which were then pooled. [a]Control groups: NS0, no substrate addition at 0 h; NS24, no substrate for 24 h. Prebiotic groups in a 24 h *in vitro* anaerobic batch fecal fermentation system: FS24, fructooligosaccharide; IN24, inulin; GS24, galactooligosaccharide; XS24, xylooligosaccharide; FL24, 2'-fucosyllactose.

**Supplementary Table 8.** Inter-study validation (ISV) average balanced accuracy according to regularization parameter $C$.

| **Inverse regularization strength parameter ($C$)** | **ISV balanced accuracy (%)** |
|---|---|
| 0.003 | 70.89 |
| 0.01 | 74.25 |
| 0.03 | 75.76[†] |
| 0.1 | 74.14 |
| 0.3 | 72.72 |
| 1.0 | 71.39 |
| 3.0 | 69.59 |

[†]Highest ISV average balanced accuracy with $C$ = 0.03.
